# Transcriptomic analysis of whole staged ovarian follicles reveals stage-specific folliculogenesis signatures in mice

**DOI:** 10.1101/2025.03.20.644278

**Authors:** Hannah VanBenschoten, Yu-Ying Chen, Olha Kholod, Daniela D Russo, Alex K Shalek, Francesca E Duncan, Teresa K Woodruff, Brittany A Goods

## Abstract

Activation and maturation of ovarian follicles are essential for female reproduction, yet the underlying molecular and transcriptional mechanisms that govern these processes remain poorly understood. In this study, we used single follicle RNA-sequencing (RNA-seq) to identify transcriptional signatures of staged ovarian follicles, from primordial to secondary stages, to uncover the genes and pathways involved in early folliculogenesis. Our findings demonstrate that primordial follicles are transcriptionally distinct from growing follicles, with enrichment in DNA integrity and RNA processing pathways, which may play a role in preserving oocyte genomic stability and cell state during dormancy. Additionally, our analysis reveals minimal transcriptomic differences between primary and secondary follicles using traditional differential expression analysis. To better distinguish growing follicle stages, we introduce unsupervised approaches, including discrete-variable predictors of follicle stage and weighted gene co-expression analysis. We identified pathways involved in DNA integrity, meiotic arrest, and cellular metabolism that drive the transition from dormant to active follicle states, as well as pathways related to cellular growth, ECM organization, and biosynthesis in growing follicle stages. Our study offers novel insights into the molecular mechanisms governing early follicle activation and growth, providing a foundation for future research with applications in reproductive biology, contraception, and fertility preservation.

**Author Summary:** The development of ovarian follicles is essential for female fertility, but the molecular signals that control their growth remain unclear. In this study, we used advanced gene sequencing techniques to analyze the genetic activity of individual ovarian follicles at different stages of early development. We found that dormant follicles have unique gene expression patterns that help protect the genetic material of the egg and maintain their inactive state. In contrast, follicles that have begun to grow show increased activity in genes related to cell growth, communication, and structural changes. Interestingly, we observed that early growing follicles are more similar to each other than previously thought, prompting us to apply new analytical methods to better distinguish their developmental stages. Our findings highlight key biological pathways that regulate the transition from dormant to active follicles and uncover new genes that may play a role in this process. Understanding these mechanisms provides valuable insights into ovarian biology and could inform future research on fertility treatments, contraception, and reproductive health.

## Introduction

The ovarian follicle is the fundamental functional unit of the ovary, consisting of the oocyte and its surrounding somatic cells. Folliculogenesis, the growth and maturation process of follicles, involves a series of critical transitions that begin with the activation of dormant primordial follicles into a state of active growth. During this transition, the oocyte undergoes significant transcriptional and translational upregulation, promoting cell growth and preparing for future developmental stages and eventual meiotic resumption (1). Concurrently, the surrounding somatic cells, which are granulosa cell precursors called pre-granulosa, take on a mature granulosa cell phenotype, exit their quiescent state, differentiate from squamous into a cuboidal granulosa cell phenotype, and resume mitotic cell division, thereby initiating the development of a larger, multilayered follicle structure (2). Developing oocytes and granulosa cells undergo symbiotic processes of maturation via bi-directional communication, endocrine signaling, and homeostatic microenvironment regulation (3,4). Ultimately, the process initiated at primordial follicle activation may culminate in the development of a fully mature follicle capable of ovulating a fertilizable oocyte (5).

Regulation of this activation process is vital for normal reproductive function, as well as for the potential preservation of fertility. The ovarian reserve is composed of a finite pool of primordial follicles that dictates reproductive lifespan and is progressively depleted over time, resulting in diminished fertility and eventual menopause. Elucidating the precise molecular cues that control activation and maturation of primordial follicles is critical to understanding key processes in reproductive biology and towards translational aims such as fertility preservation and contraception (6). Several classical signaling pathways involved in follicle activation have been described, including Wnt, insulin, Notch, and Hedgehog pathways, which all interact with the forkhead box O (FOXO) transcription factor family and the phosphatidylinositol 3-kinase/ serine/threonine kinase (P13K/AKT) pathway (7). Moreover, several key oocyte-specific genes such as *Nobox* (8), *Figla*, and *Sohlh1* (9), as well as endocrine factors such as anti-Müllerian hormone (*Amh*) (10,11) are known to play a role in follicle activation. However, we still do not fully understand the overarching systemic or temporal landscape that dictates where, how, and when dormant follicles are selected for activation, and how factors like follicle or oocyte size play into this process. Thus, despite the critical role of follicle activation in reproductive health, the complex signals that initiate and orchestrate this process remain poorly understood.

Efforts to elucidate these regulatory pathways have been hampered by the limitations of available transcriptional data, particularly regarding stage-specific follicular cells. Most studies to date have been performed on bulk RNA from whole ovaries or dissociated tissues, which fail to preserve a mapping between individual follicles and the resulting data (12,13). This approach obscures the distinct contributions of the oocyte and somatic cells at different stages of follicle development. Studies leveraging single-cell RNA sequencing to explore follicle dynamics have lacked the resolution to distinguish transcriptional changes specific to different follicle stages, such as the transition from primordial to primary and secondary follicles (14–16). As such, we previously reported single-cell transcriptomic data from somatic and germ cells dissociated from isolated staged follicles to identify drivers of follicle activation (17). This study revealed stage-specific transcriptomic profiles, showing, for example, that key metabolic and signaling pathways in the oocyte, such as the P13K/AKT pathway, are significantly upregulated during the transition from primordial to primary stages, supporting an active role in growth initiation. These findings underscore the value of RNA-seq in mapping specific transcriptional landscapes in follicular development.

Building on this staging protocol, we applied it to whole-follicle transcriptomics in the present study to characterize broader transcriptional dynamics. To date, consensus transcriptional signatures of intact primordial, primary, and secondary follicles have not been defined, and thus the whole-follicle transcriptome encompassing relative contributions of several dynamic cell types at precise development stages is unknown. Therefore, we generated transcriptomic data from individual whole follicles across primordial, primary, and secondary stages. By profiling individual follicles with morphologically-defined staging methods, we preserved the structural context of the oocyte and its surrounding somatic cells without the need for dissociation. We also demonstrate multiple computational approaches to define whole-follicle stage-specific transcriptional signatures, and each revealed genes and pathways involved in primordial follicle activation and growth. Our study captures stage-specific gene expression patterns, revealing a suite of genes and pathways that are associated with each stage.

## Results

### Staged follicles express canonical folliculogenesis markers

To characterize transcriptomic changes during follicle activation, we isolated primordial through secondary stage follicles from postnatal day 6 CD-1 mice by iterative enzymatic and mechanical digestions in accordance with previously published methods (Fig 1A) (17). Follicle stages were classified as primordial with a full layer of squamous pre-granulosa cells, primary with a full layer of cuboidal granulosa cells, and secondary with two layers of granulosa cells (Fig 1B-D) (18,19). We generated bulk transcriptomic profiles of isolated follicles to determine gene signatures that encompass the relative contributions of primary cell types (oocyte, pre-granulosa, granulosa) at each stage of follicle development. We performed Smart-Seq2 sequencing on 147 individual whole follicles at the primordial (n=63), primary (n=42), and secondary stages (n=34). Variations in sequencing depth and messenger RNA quantity can significantly affect the repeatability and biologic interpretability of bulk RNA sequencing data (20). As such, we applied the most stringent quality control filtering measures that significantly reduced the number of included follicle samples. We filtered out samples with greater than 40% mapping rate to rRNA and less than 8 million total reads. This resulted in a final dataset with 9 primordial, 6 primary, and 10 secondary follicles of high quality that were included in batch correction and downstream analysis. We used ComBat-seq (21) to correct for batch effects, which resulted in minimal intra-batch associations among samples (Fig S1).

**Figure 1.**
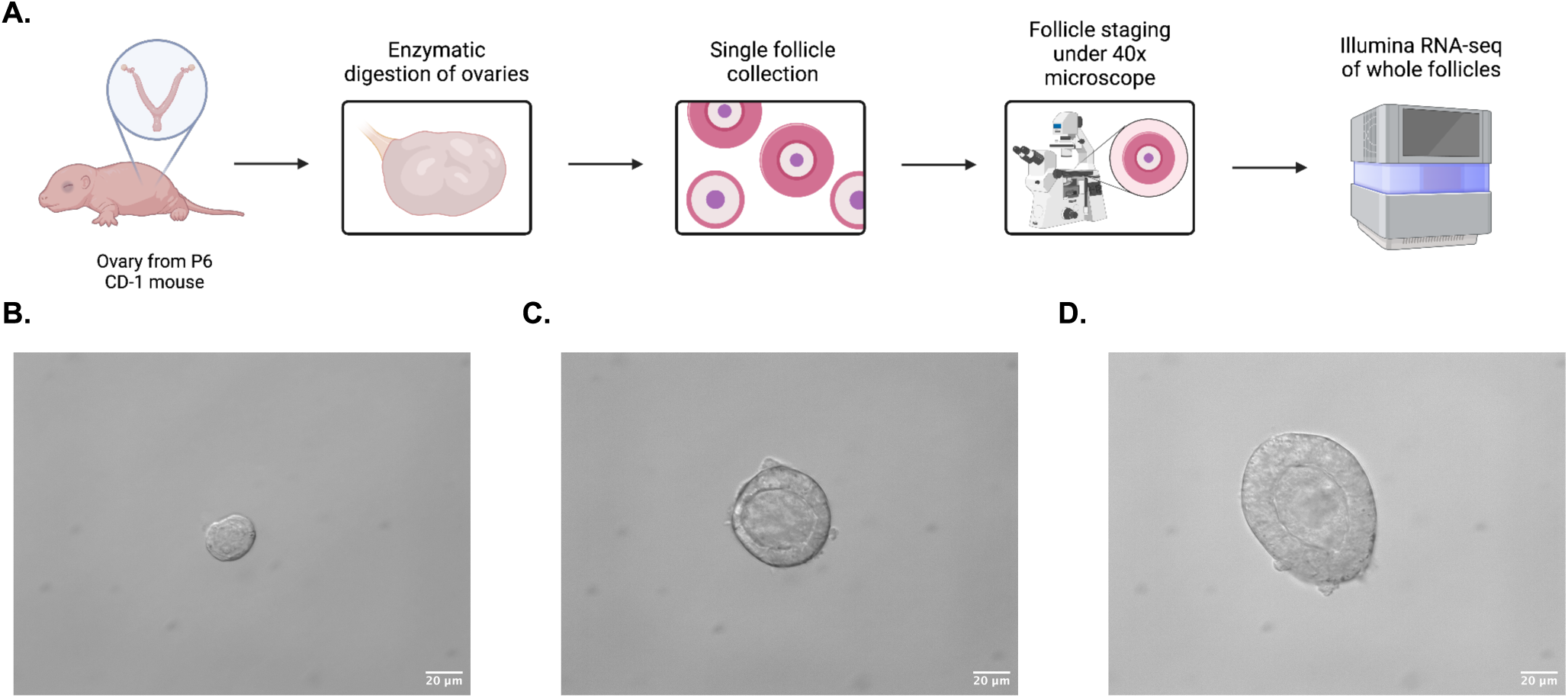
Illustration of follicle isolation and staging protocol. (A) Workflow of follicle isolation and staging. Ovaries were dissected from postnatal day 6 (P6) CD-1 mice and digested enzymatically with liberase and DNase I along with mechanical digestion. Isolated follicles were collected and staged under 40X brightfield microscopy based on defined morphological features. Single follicles were processed for bulk RNA sequencing and downstream data analysis. (B) Primordial follicles were identified by a single layer of squamous pre-granulosa cells surrounding an oocyte. (C) Primary follicles were identified by a full layer of cuboidal granulosa cells surrounding an oocyte. (D) Secondary follicles were identified by the presence of two layers of somatic cells surrounding an oocyte. Figure 1A created with BioRender. Scale bar = 20 μm.

After filtering and batch correction, a clear separation between primordial and primary and secondary follicles was apparent upon principal component analysis (PCA) (Fig 2A). While primordial follicles clustered distinctly, primary and secondary follicles were nondistinctive from each other along PC1, PC2, and every subsequent principal component visualized (Fig S2). Several known and novel follicle development genes were involved in differentiating primordial from growing follicles on PC1 (Fig 2B). Primordial follicle separation on PC1 was driven by oocyte-specific transcription factors *Sohlh1* and *Figla,* which are known to regulate the transition from primordial to primary follicles and regulate genes involved in primordial follicle survival, respectively (22). Genes involved in meiotic processes, such as *Sycp3* and *Syce1* which are involved in the synaptonemal complex of meiotic prophase, as well as *Mael* and *Meiosin*, also influenced primordial follicle clustering on PC1 (23). Genes previously not known to regulate primordial follicle survival or activation include *Camk1g*, *Ap3b2*, and *Epcam*, a cell adhesion molecule that may be linked to pre-granulosa cell transition to granulosa cells (24). Several canonical granulosa cell-associated genes drove primary and secondary follicle separation in PC1 (Fig 2B), such as *Amh*, *Fgf8*, *Hsd3b1*, and *Inhbb*, all of which are involved in granulosa cell survival, proliferation, and function (5). Genes with previous unknown roles in follicle development that drove clustering of growing follicles include *Vrtn*, *Fbln2*, *Pabpc1l*, *Scube1*, *Vcan*, and *Acsbg1*. These genes are broadly involved in ECM maintenance and remodeling, basement membrane formation and integrity, angiogenesis and vascular development, and lipid metabolism (25–30).

**Figure 2.**
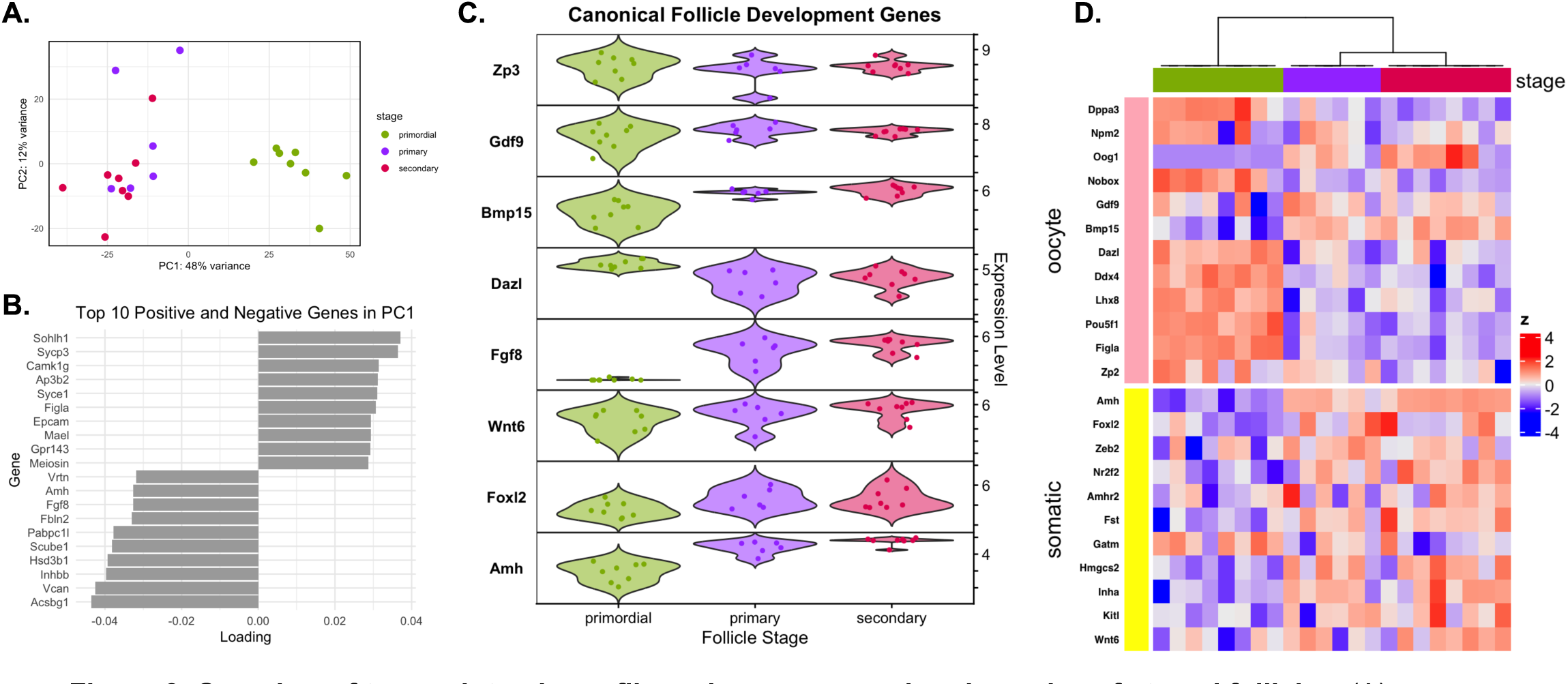
Overview of transcriptomic profile and gene expression dynamics of staged follicles. (A) Principal component analysis (PCA) of whole follicles (8 primordial, 6 primary, 8 secondary) after collection and computational filtering for transcript quality. Follicles colored by stage showed distinct separation between primordial follicles and primary and secondary follicles along PC1. (B) Top 10 positive and negative weighted genes that drove principal component 1 separation. Genes with positive loadings correspond to primordial follicles and genes with negative loadings correspond to primary and secondary follicles. (C) Expression level (logTPM of count matrix from DESeq2) of canonical follicle development genes across follicle stages plotted on violin plots. (D) Heatmap of expression (z-score) of known oocyte and somatic-cell specific genes curated from single cell ovarian follicle dataset plotted across follicle stages.

To verify the expression and dynamics of genes with known involvement in follicle development, we plotted the normalized expression of canonical follicle development genes across follicle stages (Fig 2C). Canonical oocyte development marker *Zp3* was expressed broadly across all follicle stages. *Gdf9* has previously been reported to only be expressed after the primary stage (17) and is believed to play a role in the transition from primary to secondary follicles (31). In coordination with upregulated *Bmp15* expression at the primary stage, *Gdf9* regulates oocyte-granulosa cell communication and influences the sensitivity of granulosa cells to follicle stimulating hormone resulting in their proliferation (32); both genes play a role in reducing early follicular luteinization by suppressing progesterone production. While *Gdf9* and, to a lesser extent, *Bmp15*, are expressed heterogeneously in primordial follicles, their expression increases consistently upon primary follicle transition as expected. Oocyte-specific gene *Dazl* was highly expressed in primordial follicles with lower, more varied expression across primary and secondary follicle stages. Conversely, *Fgf8* expression was not detected in primordial follicles and increased dramatically in primary and secondary follicles, as expected (33). Somatic cell specific genes *Wnt6*, *Foxl2*, and *Amh* were detected across follicle stages with relatively increased expression at later stages, particularly for *Amh*.

A larger set of known oocyte and somatic cell-specific genes was also plotted to illustrate broad changes in the relative contributions of oocyte and somatic cell signals to the whole staged follicle transcriptome (Fig 2D). While genes with known expression dynamics in growing follicles (*Gdf9*, *Bmp15*, and *Oog1*) showed increased expression in primary and secondary follicles, oocyte-specific genes largely showed greater relative expression in primordial follicles. These findings validate known differences in transcriptomic characteristics between follicle stages, suggesting our dataset comprised of high-quality follicle data post stringent filtering that can be used for finding new signatures of folliculogenesis.

### Differential expression across follicle stage identifies gene expression signatures between primordial and growing follicles

We next performed differential expression analysis to compare the transcriptomes of primordial, primary, and secondary follicles. We identified 1,244 differentially expressed genes (DEGs) (log fold change >1 & P_adj_ < 0.05) across all comparisons (Fig 3A, Fig S3). Primordial follicles expressed 503 upregulated and 737 downregulated genes in comparison to both primary and secondary follicles. Conversely, there were only 4 DEGs identified between primary and secondary follicles, with all 4 upregulated in primary follicles. Novel DEGs with previously unreported involvement in follicle development were also identified (Fig 3B). Genes with upregulated expression in primordial follicles include *Stpg1*, *Camk1g*, *Tex13b*, *Card10*, *Ttc22*, and *Colec11*. Genes with both known and novel roles in follicle development were upregulated in primary and secondary follicles in comparison to primordial follicles, including *Shisa8*, *Eya2*, *Mrap*, *Fbxw20*, *Myo7a* and *Ank1*. Besides *Shisa8*, which modulates Wnt and FGF signaling pathways that have known roles in folliculogenesis at the primary and secondary stages (7), these genes have unstudied roles in follicle development.

**Figure 3.**
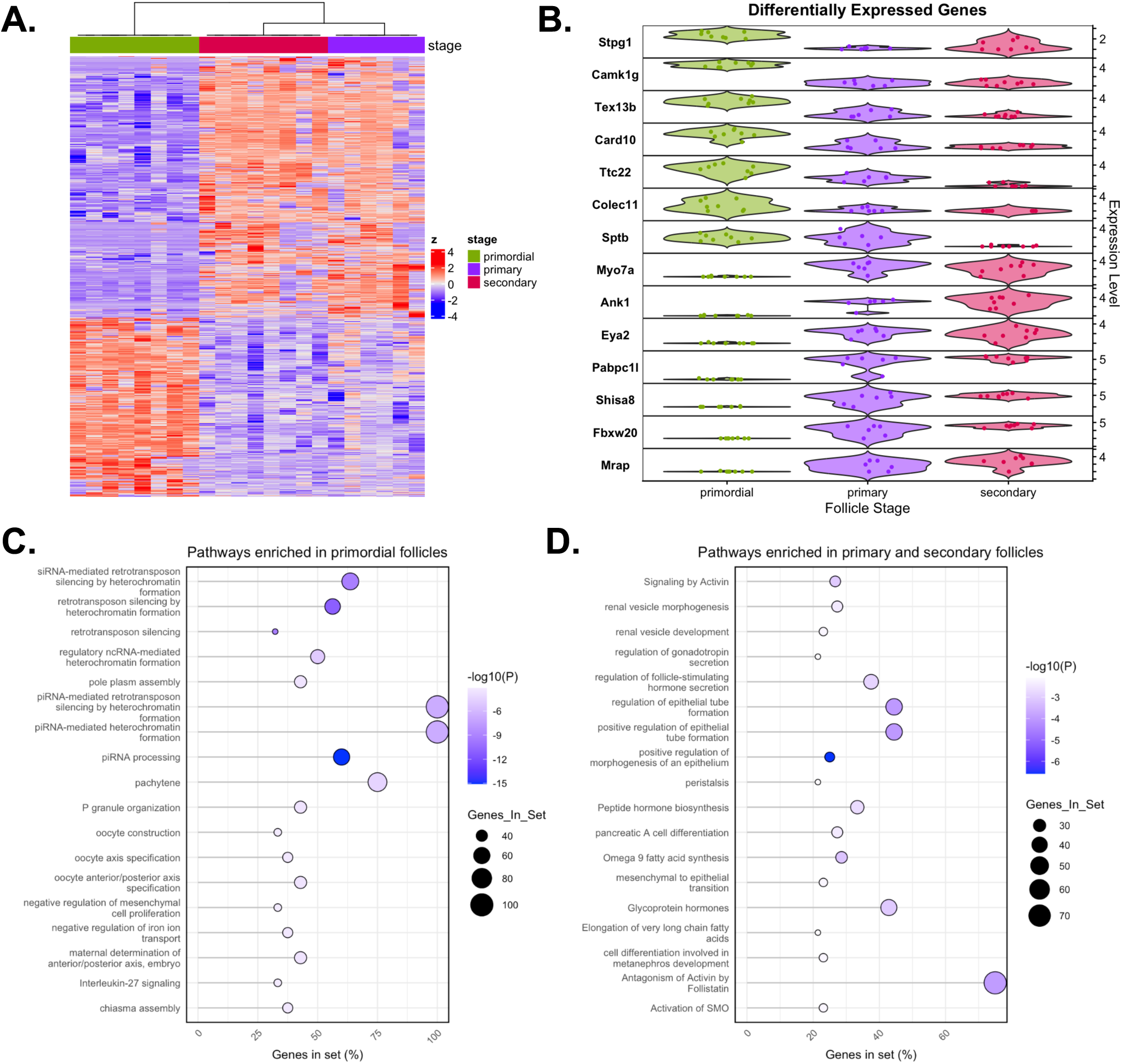
Differential gene expression across follicle stages reveals distinct gene signature between primordial and primary/secondary follicles. (A) Heatmap of the union of significantly differentially expressed genes (Log fold change > 1 and p_adj_ < 0.01) across primordial, primary, and secondary follicles. (B) Violin plots of expression (normalized counts from DESeq2 object) of top significantly differentially expressed genes across follicle stages. Functional enrichment analysis dotplots of upregulated genes in (C) primordial follicles and (D) primary and secondary follicles (union). Gene set enrichment annotations were generated in Metascape and represent pathways and functions from GO Biological Processes, KEGG, WikiPathways, and Reactome. Terms in which greater than 30% (C) and 20% (D) of the total genes in the set were expressed in upregulated gene lists are plotted on the x-axis and visualized by circle size. Circle fill color represents the significance determined by p-value (-Log10(P)).

To understand the biological functions driving differential gene expression at primordial and growing follicle stages, we conducted gene set enrichment analysis using Metascape (34). Functional enrichment analysis dot plots of upregulated genes in primordial follicles (Fig 3C) and primary and secondary follicles (union of DEGs) (Fig 3D) were generated from GO Biological Processes, KEGG, WikiPathways, and Reactome databases. There is significant representation of epigenetic maintenance and genome integrity processes via retrotransposon silencing in the primordial phase, as well as meiotic cell processes that relate to meiotic prophase including pachytene. In primordial follicles, piRNA processing and retrotransposon silencing are critical for maintaining genomic stability, protecting oocytes from transposable element activity (35). Pathways involved in oocyte axis specification, pole plasm, and chiasma assembly highlight early activity in oocyte polarity establishment, which is typically thought to occur after meiotic resumption in the preantral follicle in mice (36). Epithelial regulation and morphogenesis, hormone responsiveness and secretion, and activin antagonism by follistatin are major pathways represented in primary and secondary follicles.

### Follicle and oocyte size correlate with stage- and cell-specific DEGs

We performed correlation analyses between follicle and oocyte size and gene expression. Violin plots comparing follicle and oocyte sizes across primordial, primary, and secondary follicles show that both follicle and oocyte size significantly increase as follicles mature, as expected (Fig 4A). These size distributions reflect the distinct growth phases associated with each follicle stage that correlate with morphological changes used in their characterization. Using EdgeR, we identified genes that are significantly correlated (|Log fold change| > 1.5 and p_adj_ < 0.05) with follicle size and oocyte size (Fig 4B). There were 906 shared genes that correlated with both sizes, 342 genes that correlated exclusively with follicle size, and 227 genes that correlated exclusively with oocyte size. GSEA analysis of follicle size specific genes revealed a high correlation of follicle size with the OAP-1-OSP/claudin-11-ITGB1 complex (Fig S4). Platelet activating factor biosynthetic and metabolic processes were overrepresented in oocyte size specific pathways (Fig S4). Overall, many genes are broadly involved in both overall follicular and oocyte development during folliculogenesis, however, a substantial portion of genes are exclusively correlated to one size parameter.

**Figure 4.**
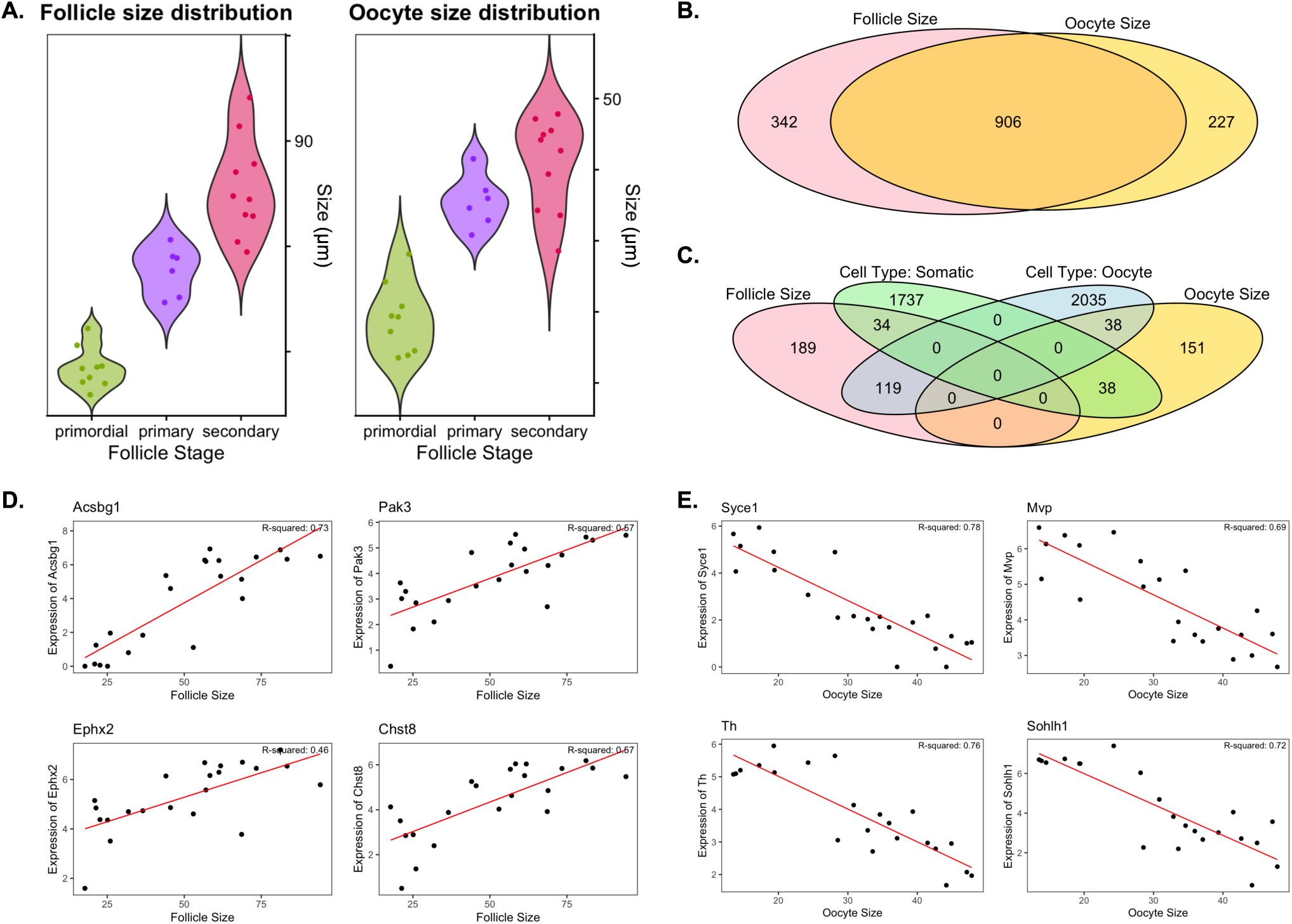
Follicle and oocyte size correlate with stage-specific DEGs. (A) Violin plots of measured follicle size (left) and oocyte size (right) across primordial, primary, and secondary follicles. (B) Venn diagram of genes that were significantly correlated (Log fold change > 1.5 and p_adj_ < 0.05) with follicle size (pink) and oocyte size (yellow), with overlapping genes represented by the middle ellipse (orange). Numerical labels represent the total number of genes in each set. (C) Four-set Venn diagram of genes that are significantly correlated to follicle size (pink) and oocyte size (yellow), as well as somatic cell (green) and oocyte (blue) specific genes as determined by differential expression analysis of a previously published single cell P6 mouse ovarian follicle dataset. Intersections and numeric labels represent the total number of shared genes between each group. Normalized expression (Log_2_TPM) of top 4 genes that most closely correlate with (D) follicle size as a function of measured follicle size in µm and (E) oocyte size as a function of measured oocyte size in µm. Individual datapoint represent each follicle sample and red lines represent linear regression with corresponding R-squared values.

To explore the cell-specificity of genes correlated with follicle and oocyte size, we integrated our analysis with differentially expressed genes from our previously published single-cell ovarian follicle dataset that contains somatic cells and oocytes across primordial, primary, and secondary stages (17). We compared genes upregulated in oocytes and in somatic cells (identified using DESeq2) with genes that were uniquely correlated to either follicle or oocyte size (Fig 4C). Notably, 1,809 genes were upregulated in somatic cells (Log fold change > 1.5 and p_adj_ < 0.05) and of that total, 34 genes overlapped with follicle-size correlated genes while 38 gene overlapped with oocyte size correlated genes. By contrast, 2,192 genes were upregulated in oocytes (Log fold change > 1.5 and p_adj_ < 0.05), wherein 38 overlapped with oocyte size correlated genes and 119 overlapped with follicle size correlated genes. Most of the follicle size correlated somatic cell genes were positively correlate with follicle size (27 positive versus 7 negative); similarly, the majority of oocyte size correlated - oocyte genes were positively correlated with oocyte size (26 positive versus 12 negative). Interestingly, only one follicle-size-oocyte-specific gene had a positive correlation with follicle size (*Prr18*), while 118 genes were negatively correlated with follicle size. Conversely, only one oocyte-size-somatic-cell-specific gene had a negative correlation with oocyte size (*Sptas2l*), while 37 genes had a positive correlation. Gene ontology terms that correlate with follicle and oocyte size that are somatic cell or oocyte specific were identified (Fig S5).

We next plotted the normalized expression of the top 4 most highly correlated genes to follicle size that were somatic-cell specific (Fig 4D) and oocyte size that were oocyte-specific (Fig 4E). Somatic cell-specific genes *Acsbg1*, *Pak3*, *Ephx2*, and *Chst8* positively correlated with increasing follicle size. These genes share roles in lipid metabolism (*Acsbg1* and *Ephx2*) (26,37) and in cytoskeletal remodeling, signaling, and extracellular matrix dynamics (*Pak3*) (38), thus they likely contribute to the various processes required for follicular growth and maturation, supporting somatic cell expansion and restructuring that occurs in primary and secondary follicle stages. *Chst8* belongs to the sulfotransferase 2 family and catalyzes the sulfination of luteinizing hormone, which is critical for regulating the timing of sexual maturation (39). In contrast, oocyte-specific genes *Syce1*, *Mvp*, *Th*, and *Sohlh1* negatively correlated with increasing oocyte size. *Syce1* has known involvement in the synaptonemal complex of meiosis and is important for maintaining the chromatin integrity in oocytes during prophase I meiotic arrest (23). Similarly, *Mvp* encodes major vault protein that is involved in forming vault complexes, which are characteristically implicated in metastasis of many cancers and have an unknown role in oocytes of primordial follicles (40,41). *Sohlh1* has a known role in early oogenesis and folliculogenesis in the primordial phase. Interestingly, it was previously reported that oocytes exhibit no expression of *Th*, a tyrosine hydroxylase associated with catecholamine biosynthesis, thus its expression and role in oocytes remains to be further elucidated (42). These findings point to known and novel stage-specific gene regulation that is correlated with growth and maturation processes in both the somatic and germ-cell compartments of a growing follicle. This analysis reveals key genes whose expression is tightly linked to oocyte and follicle size and provides an additional method of identify specific transcriptomic programs active during the transition from primordial to secondary stages.

### Unsupervised gene expression and pathway analysis methods reveals modules associated with follicle stage

We next determined if we could identify modules of co-regulated genes that may drive folliculogenesis. We used weighted gene co-expression network analysis (WGCNA) (43) to identify gene modules associated with follicle stage-specific transcriptional profiles. The module-condition association heatmap (Fig 5A), where each row corresponds to a unique gene module and each column to a specific follicle stage, shows gene modules that correspond highly to each follicle stage. Pearson correlation coefficients between each module eigengene and the follicle stage are represented within the cells, with p-values provided in parentheses. Three modules showed the highest stage-specific associations: the brown module was strongly correlated with primordial follicles (r = 0.76, p = 9e-05), while the grey and black modules were most strongly associated with primary and secondary follicles, respectively (r = 0.68, p = 2e-06 and r = 0.63, p = 0.005). Heatmaps illustrate gene expression within these top stage-correlated modules (Fig 5B). Importantly, individual follicles and their unique gene co-expression patterns had a significant impact on these module-condition associations. For example, in the brown module, one primordial follicle exhibited high expression of nearly all genes in the module, while the remaining primordial follicles displayed high expression of only half or fewer genes. This suggests that we identified many modules that describe underlying heterogeneity within developmental stage.

**Figure 5.**
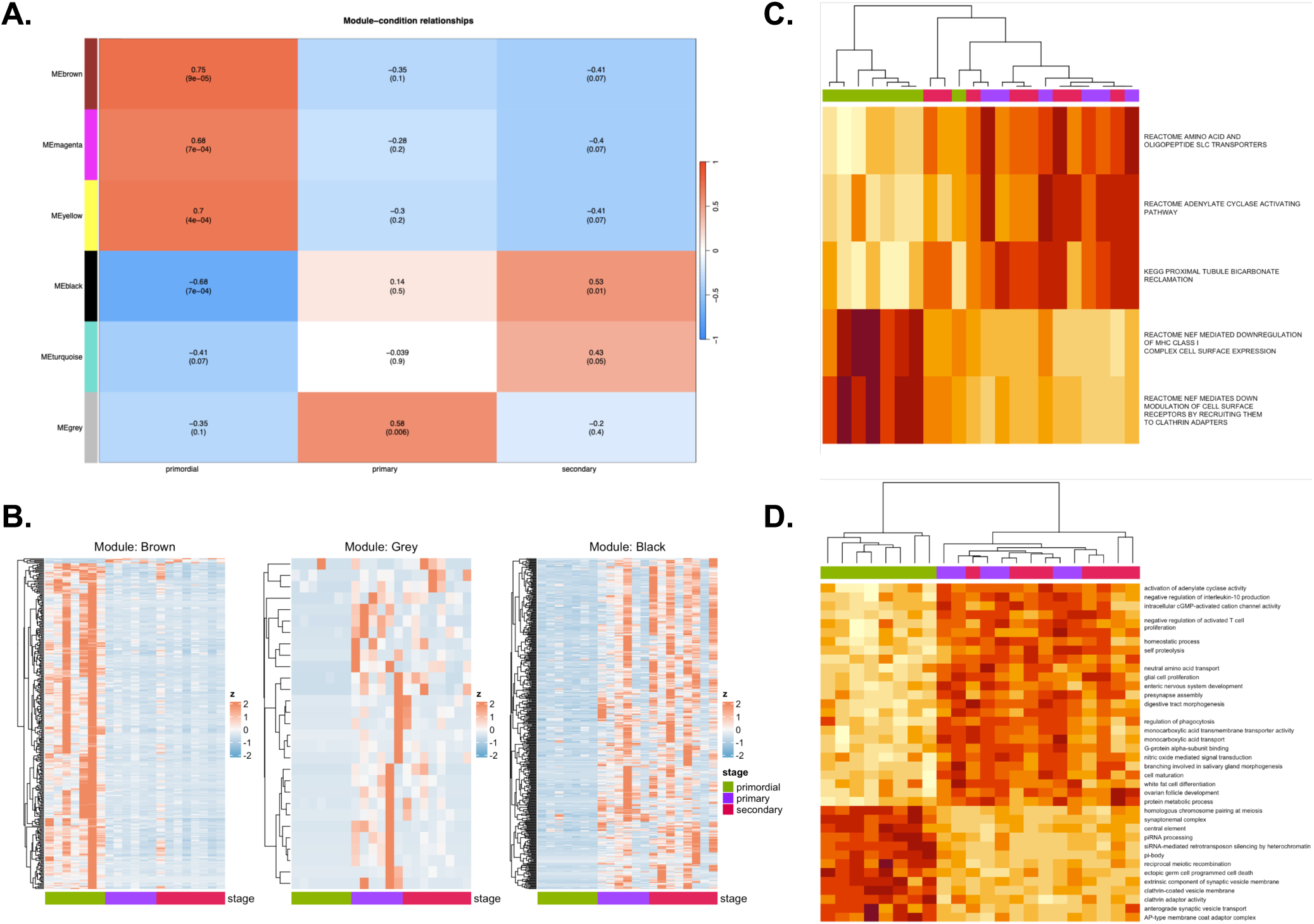
Weighted correlation network analysis (WGCNA) reveals gene modules driven by follicle heterogeneity. (A) Module-condition association heatmap from WGCNA analysis wherein each row corresponds to a unique gene module and each column corresponds to follicle stage as labelled. Cells contain Pearson correlation coefficient between module eigengene and follicle stage (condition) with corresponding p-value in parentheses. (B) Heatmaps of gene expression within three modules that are most closely correlated to each follicle stage (e.g. Brown = primordial; grey = primary; black = secondary). (C) Heatmap of differential pathway expression (from KEGG, Reactome, and Biocarta) among staged follicle samples as determined by Gene Set Variation Analysis (GSVA). (D) Heatmap of differentially expressed GO Biological Process, Molecular Function, and Cellular Component annotation sets as determined by GSVA. Columns represent samples clustered based on their calculated pairwise Spearman correlation coefficients and rows represent pathway or gene set analysis annotations note databases. Cell color represents the row-normalized expression levels of differentially expressed pathways or gene sets.

We further characterized pathways active in individual staged follicles using gene set variation analysis (GSVA) (44), which allowed for unsupervised, gene set concordance-based analysis of enriched pathways (Fig 5C). Primordial follicles exhibited enriched expression in pathways associated with cell surface expression. Primary and secondary follicles showed strong association with tubule bicarbonate reclamation; a pH mediation pathway typically ascribed to the kidneys, as well as hormone responsiveness (adenylate cyclase activating pathway) and metabolic activity via nutrient transport (amino acid and oligopeptide SLC transporters). Further GSVA analysis of Gene Ontology (GO) terms and cellular components (Fig 5D) revealed distinct biological processes and molecular functions driving each stage of folliculogenesis. Primordial follicles exhibited enriched GO terms associated with genome stability and cell-state maintenance (pi-RNA processing, si-RNA-mediated retrotransposon silencing), in accordance with prior GSEA analysis on DEGs (Fig 3C). Clathrin adaptor activity and vesicle transport terms appeared in both pathway and gene ontology-based GSVA of primordial follicles. Primary and secondary follicles showed enrichment in cellular processes that largely confirmed pathways results from GSVA (Fig 5C), namely meiotic arrest by cAMP (adenylate cyclase activity, intracellular cGMP-activated cation channel activity), pH regulation (monocarboxylic acid transport), and amino acid transport. Cell proliferation processes and morphogenesis were also largely represented. Overall, this gene set concordance-based analysis underscores the transcriptional complexity underlying follicle development, pinpointing specific gene modules and pathways that contribute to follicular stage progression and heterogeneity.

## Discussion

Here, we identified transcriptional signatures of staged whole ovarian follicles to uncover pathways and gene targets involved in early folliculogenesis. Through bulk transcriptomic analysis, we successfully captured the dynamic molecular landscape of follicle development, from primordial to secondary stages. Our findings offer new insights into the gene expression patterns that characterize each stage of folliculogenesis across stage and as a function of size. Our results demonstrate that primordial follicles, in contrast to growing follicles, are characterized by distinct transcriptional profiles that reflect their quiescent state and need to maintain genomic stability (Fig 6). In contrast to primary and secondary follicles, primordial follicles dominantly express known oocyte-specific genes. This is likely because oocytes are a relatively larger component of primordial follicles compared to primary or secondary follicles, having only a single layer or pre-granulosa cells. Conversely, somatic cell specific genes generally show greater relative expression in primary and secondary stages due to granulosa cell differentiation and expansion. The strong enrichment of DNA integrity maintenance processes, including piRNA processing and retrotransposon silencing by heterochromatin, is consistent with knowledge of the role of epigenetic regulation in maintaining cell-type identity and quiescence in dormant primordial follicles (45). This includes mechanisms to regulate and silence transposable elements, which can be reactivated during the long dormancy period of primordial follicles and play a role in oocyte degradation during reproductive aging (46,47). We note the enrichment of processes associated with the diplotene phase of meiotic prophase I in Gene Set Enrichment of primordial follicles (synaptonemal complex, homologous chromosome pairing, chiasmata, etc.) (Fig 6) (48). This suggest that genes involved in diplotene (e.g. *Cdc25b*, *Sycp1*, *Sycp3*) are actively transcribed in primordial follicles during meiotic arrest. There is some disagreement in our findings with oocyte meiotic progression, specifically oocyte axis specification as identified GSEA of primordial follicles. Oocyte polarization is initiated upon germinal vesicle breakdown, which triggers metaphase 1 in antral follicles; thus, it is curious to note axis specification in early-stage follicle development (49). With further validation, staged follicle transcriptomics could be used to probe the synchronicity of follicular development with oocyte meiotic progression. The transition from primordial to primary follicles is marked by significant transcriptional changes, particularly the upregulation of genes involved in granulosa cell proliferation, extracellular matrix (ECM) remodeling, and oocyte maturation (Fig 6). Pathway analysis confirms known processes involved follicle activation in primary follicles. SMO is the smoothened transmembrane signaling protein whose activation is a central component of the Hedgehog signaling pathway, which is known to become activated in granulosa cells at the primary follicle stage (50). Similarly, activin, a member of the TGF-β superfamily, is known to play a role in growth and differentiation of granulosa cells as well as enhancing follicle responsiveness to FSH, which is essential for follicular transition to advanced stages (51–53).

**Figure 6.**
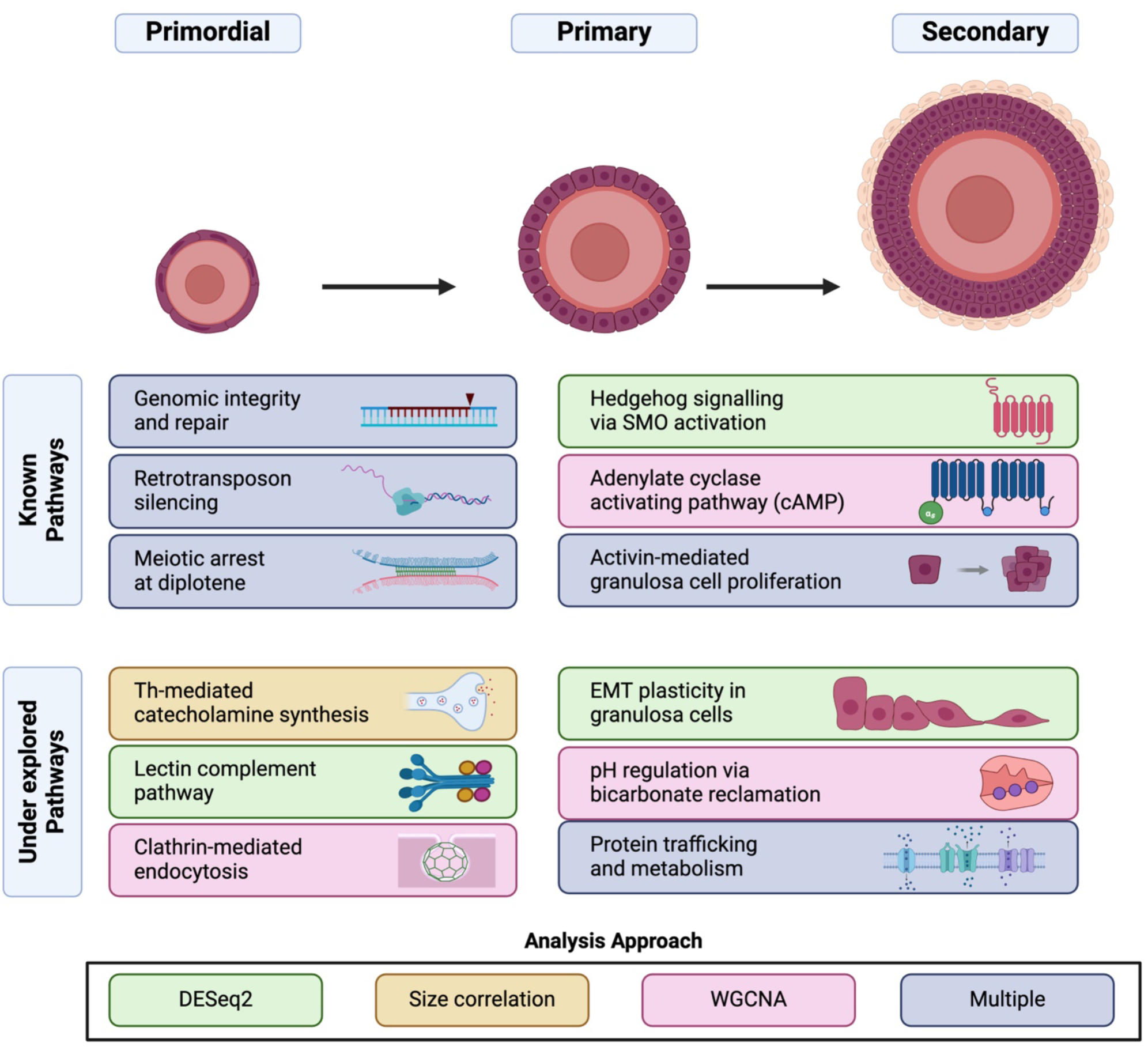
Summary of key findings. Different stages of early folliculogenesis exhibit distinct transcriptomic and phenotypic states. Primordial follicles are known to undergo meiotic arrest in the diplotene phase of prophase I, thus pathways involved in genetic integrity and repair and retrotransposon silencing are concordant with the maintenance of cell-state in this phase. We identified several underexplored pathways in primordial follicles, including the capacity for oocyte-intrinsic Th-mediated catecholamine synthesis, which is involved in follicular innervation, the role of the lectin complement pathway in primordial follicle patterning, and clathrin-mediated endocytosis during steroid trafficking. In contrast with primordial follicles, primary and secondary follicles shared similar transcriptomic signatures; genes activated in primary follicles were persistently expressed in secondary follicles. Known pathways such as hedgehog signaling, cAMP, and activin-mediated granulosa cell proliferation were reflected in our data, as well as under explored pathways such as EMT plasticity in granulosa cells, intercellular pH regulation, and protein trafficking and metabolism.

Identification of novel genes that have not been previously linked to follicle activation and growth suggests new avenues of research into the molecular mechanisms regulating these processes. In primordial follicles, several genes with unknown roles in folliculogenesis were upregulated. *Stpg1* is a protein encoding gene that is predominantly expressed in the testes and is believed to play a role in spermatogenesis and the structure and function of the sperm tail, but it’s role in ovarian biology has not been reported (54). Similarly, *Tex13b* is predominately expressed in the testis with no characterized role in ovarian development, though *Tex14* is known to be involved in intercellular bridge formation between embryonic germ cell (55,56). *Camk1g* is a member of the CaMK kinase family with a known role in calcium signaling pathways involved in cell communication and development; while other CaMK protein subfamilies have a known role in oocyte and follicular competence, the role of CaMK1 in folliculogenesis has not been explored (57). Indeed, prior studies have focused solely on the role of *Camk2* in oocyte maturation (57) and it was recently demonstrated that *Camk2* inhibition prevents primordial follicle activation (58). The upregulation of *Camk1* in primordial follicles suggests that a more robust understanding of protein kinases involved in calcium signaling during follicle activation is warranted. *Card10* is a caspase recruitment scaffolding protein involved in NF-kB signal transduction leading to apoptosis with an undefined role in follicle development (59). Another protein-protein interaction gene, *Ttc22*, is not well characterized and has no evidence which links it to ovarian follicle development. Finally, *Colec11* encodes C-type lectins involved in the lectin complement pathway, which plays a role in neural crest cell migration and organ patterning during development; thus, its role in primordial follicles, while undefined, could relate to primordial follicle formation and patterning (Fig 6) (60). Various components of the complement pathway have been identified in human follicular fluid and the lectin pathway is known to stimulate cell migration, thus it may play a role in recruitment of mesenchymal cells from the ovarian stroma to stimulate follicle activation (5,61,62). *Sptb* is one of four genes with differential expression between primary and secondary follicles, wherein it is upregulated in primary follicles. *Sptb* encodes a spectrin protein involved in maintaining cell shape and membrane integrity, primarily in erythrocytes; its role in primary follicle development has not been studied (63).

In primary and secondary follicles, several novel DEGs were identified with unknown or unexplored roles in folliculogenesis. For instance, *Eya2* has been shown to play a role in epithelial-to-mesenchymal transition in reproductive tissues, particularly in the context of ovarian cancer, though its specific involvement in follicular development has not been reported (Fig 6) (64). TGF-β-mediated EMT of granulosa cells during folliculogenesis is well-characterized and suggest that a partial mesenchymal phenotype is required for normal follicle development; thus, *Eya2* may play a role in granulosa cell transition and differentiation in growing follicles (65–67). While our data suggest that *Eya2* may be involved in the transition of granulosa cells from their precursor states to functional cell types, pathway analysis also revealed the role of several genes (*Gdnf*, *Wnt4*, and *Smo)* in epithelial tube formation. It is known that despite EMT, granulosa cells develop and maintain epithelial characteristics during early follicle development, such as cuboidalization and polarization (65,68). Thus, we suggest that granulosa cell exhibit a dynamic epithelial-mesenchymal phenotype during early follicle development with involvement of previously unexplored genes, such as *Eya2*. Other novel DEGs found in primary and secondary follicles (*Mrap*, *Fbxw20*, *Myo7a* and *Ank1)* relate to steroid receptor trafficking, F-box protein-mediated proteolysis, cell movement and cell membrane integrity, respectively, and all have speculative but previously unstudied roles in follicle development (69–72). The role of novel stage-specific DEGs may be further contextualized by gene expression data with multi-omic analysis pipelines. For instance, transcriptomic data from staged ovarian follicles generated via microarray has been leveraged to infer inter- and intra-cellular metabolic networks in ovarian follicles, which revealed metabolic pathways with novel roles in folliculogenesis (73). A robust transcriptome that describes stage-specific follicle dynamics can thus enable further systems biology approaches that may elucidate underlying ovarian biology.

We employed various analytical methods beyond standard differential expression to identify genes and pathways that differentiate primordial from primary and secondary follicles and drive follicle activation and growth. Using size as predictors of follicle stage, we identified 1,475 genes total that were significantly associated with change in follicle or oocyte size, with 342 and 227 genes that correlated exclusively to either follicle or oocyte size, respectively. Interestingly, when we incorporated single-cell data from isolated granulosa cells and oocytes, we did not see exclusive overlap of follicle-size-specific genes and somatic cell genes or oocyte-size-specific genes and oocyte genes. However, we did observe that nearly all follicle-size-oocyte-specific genes were negatively correlated with follicle size, whereas nearly all oocyte-size-somatic-cell-specific genes were positively correlated with oocyte size. This may indicate directionality of cell-cell communication at different stages, wherein dominant oocyte expression in primordial follicles contributes to overall follicle dormancy, including in pre-granulosa cells, while granulosa cell signaling contributes significantly to oocyte growth in activated, growing follicles. One interesting finding from size-based transcriptomic signature analysis was the heightened expression of tyrosine hydroxylase (*Th*) in small oocytes, which suggests that oocytes themselves may have the ability to synthesize catecholamines, such as dopamine and norepinephrine (Fig 6). This challenges previous assumptions that catecholamine production in oocytes is solely reliant on external sources due to the finding that oocytes express TH coenzyme dopamine β-hydroxylase (DAB), but not TH itself. It was therefore hypothesized that intrinsic oocyte NE synthesis is facilitated by active transport of TH into oocytes by nearby ovarian catecholaminergic neurons (42). Our data suggest that oocytes could be self-amplifying sources of these neurotransmitters, potentially playing a previously unrecognized role in early follicle activation and communication with surrounding granulosa cells and further work should seek to validate this experimentally. The expression of *Th* in oocytes could open new avenues of research into how neurotransmitter signaling impacts follicle development and oocyte competence as well.

Weighted Gene Co-expression Network Analysis (WGCNA) generated distinct gene modules associated with primary and secondary follicles and further demonstrated the utility of unsupervised co-expression-based analysis in identifying distinct gene networks by follicle stage. Gene Set Variation Analysis (GSVA) was used to identify pathways associated with co-expressed gene networks, highlighting how unsupervised analysis methods can capture the complexity of follicle stage transitions. Pathways involved in DNA integrity and meiotic arrest were enriched in primordial follicles, pointing to the key biological processes that govern follicle senescence. Terms related to clathrin-mediated endocytosis were associated with primordial follicles at the pathway and gene ontology levels; the role of clathrin-adaptors in steroid trafficking in primordial granulosa cells has been suggested but not heavily explored and warrants further investigation (Fig 6) (74). Identification of the adenylate cyclase activating pathway in growing follicles, essential for maintaining meiotic arrest via cyclic AMP (cAMP) signaling, aligns with known mechanisms regulating oocyte maturation (48,75). The adenylate cyclase activating pathway involves a G protein coupled cascade that leads to increased production of cyclic AMP (cAMP) via adenylate cyclase; high intra-oocyte cAMP levels are necessary to maintain meiotic arrest until meiotic resumption is triggered by a surge in pituitary-derived luteinizing hormone (LH) (75). Indeed, concordant expression of genes in the Gs-GPR-ADYC pathway across growing follicles indicates that this pathway is activated in primary and secondary follicles (Fig 6). The amino acid and oligopeptide SLC transporter pathways involved solute carrier transporters responsible for moving amino acids and small peptide across cellular membranes; upregulation in growing follicles likely reflects the increased demand for amino acids to support rapid cellular growth, oocyte maturation, and proliferation of granulosa cells (Fig 6). This pathway may also be indicative of cell-cell communication that underpins oocyte maturation and granulosa cell expansion in growing follicles. Finally, the proximal tubule bicarbonate reclamation pathway describes bicarbonate-mediated pH regulation. While this pathway is specific for renal cells of the kidney, shared genes in our data suggest pH regulation is an important mechanism in growing follicles (Fig 6). Is has been previously shown that granulosa cells of growing follicles and cumulous-oocyte complexes (COCs) are required to maintain a homeostatic pH 7.2 in oocytes by gap-junction mediated chloride-bicarbonate exchangers (76,77). Activation of pH exchange pathways in early growing follicles has been identified specifically in follicles in the 60-65 μm diameter range (77,78), which correlates approximately to primary and secondary follicles in our dataset. Upregulation of these pathways underscores the metabolic and homeostatic demands of growing follicles as they prepare for ovulation, as well as the role of granulosa cells in maintaining a microenvironment amenable to oocyte development (78).

One of the primary limitations of our study is the small sample size. This is due to challenges in generating high quality sequencing data on isolated *in vivo* derived follicles, as shown by stringent filtering rates applied to create a very high-quality final dataset. The primary differential expression tools used here (DESeq2 and limma) can detect significantly differentially expressed genes with >90% true positivity rate and < 5% false discovery rate using a log-fold change cut off greater than one with at least six replicates. Our datasets meet these criteria, thus we are not limited in our ability to detect DEGs with stringent statistical cutoffs; rather, the main limitation of having fewer replicates is less sensitivity to identifying differentially expressed genes with subtle gene expression differences (79). Another limitation is the potential age-specific nature of the transcriptional signatures we identified. The follicles were derived from young mice, which may not fully capture the gene expression dynamics in adult ovarian follicles. This is particularly relevant for growing follicles in the secondary stage, where prior studies have demonstrated significant differences in gene expression profiles between children and adults (80). Recruitment of theca cells and pathways related to stromal signaling, which are more pronounced in adult secondary follicles and antral follicles, may not be fully represented in our dataset. Further studies with larger sample sizes and broader age ranges will be necessary to validate and extend our findings. Despite these limitations, this study advances the field by bridging high-resolution single-cell insights with bulk transcriptomic profiling, providing a complementary perspective on follicle development. These datasets provide a comprehensive reference for transcriptomic-based follicle staging, which has implications for further research in reproductive biology and translational efforts in ovarian health and fertility preservation.

## Conclusion

This study provides a comprehensive transcriptional analysis of whole staged ovarian follicles, uncovering key pathways and novel genes involved in early folliculogenesis. Our findings underscore the importance of DNA integrity and retrotransposon silencing in primordial follicles, as well as the critical roles of calcium signaling and epithelial-mesenchymal balance in growing follicles. Furthermore, our analysis highlights the heterogeneity present within follicles at each stage, suggesting varying developmental potentials even within the same morphological stage. These insights have significant implications for understanding follicle development and offer new avenues for research into fertility treatments and nonhormonal contraceptives. Future studies should focus on validating these gene targets and pathways, as well as expanding the use of spatial transcriptomics to further dissect the cellular complexity of folliculogenesis. By advancing our understanding of the molecular landscape of follicle development, this research contributes to the broader field of reproductive biology and holds potential for clinical applications in fertility and ovarian health.

## Materials and Methods

### Animals

ICR (CD-1^ⓡ^) outbred breeders were purchased from Envigo (IN, USA) and bred in house at the animal facility of Northwestern University Center for Comparative Medicine. Mice were kept in a 14-h light: 10-h darkness cycle with constant temperature and humidity, with food (Teklad 2020X, Envigo) and water given *ad libitum*. Animals were treated in accordance with the National Institutes of Health Guide for the Care and Use of Laboratory Animals. All protocols were approved by the Northwestern University Institutional Animal Care and Use Committee (IACUC #IS00010942)

### Follicle collection, measurement, and staging

For each collection, 8-12 postnatal day 6 (P6) ovaries were used for follicle isolation using the protocol as described previously (17). Briefly, ovaries were pooled and digested in 500 μL Leibovitz’s L-15 medium (Thermo Fisher Scientific, MA, USA) supplemented with 1 mg/mL poly(vinyl alcohol) (MilliporeSigma, MO, USA) (PVA, hereafter referred to as L-15/PVA), 30.8 μg/mL Liberase TM (Roche, Switzerland), and 456 U/mL DNase I (Worthington, NJ, USA) and incubated on a 37°C heated stage for 13 min. The ovaries were then rinsed in L-15/PVA before mechanical pipetting for 4 min at 25°C in 500 μL L-15/PVA supplemented with 50 μL/mL FBS (MilliporeSigma). The process was repeated for two more rounds, with the second round of digestion being 7 min L-15/Liberase incubation and 4 min mechanical pipetting. The third round was 4 min L-15/Liberase incubation and 4 min mechanical pipetting. After three rounds of digestion and pipetting, media in which ovaries were mechanically disrupted were collected and passed through a 70-μm cell strainer (Corning Inc., NY, USA) into a center well dish (Thermo Fisher Scientific). Follicles were staged and washed in three drops of 50 μL L-15/ PVA with 100 μL/mL FBS and transferred individually into each well of a 0.2 mL skirted 96-well PCR plate (Thermo Fisher Scientific) containing 10 μL buffer RLT (Qiagen, Germany) with 1% 2-mercaptaethanol (MilliporeSigma). Follicle stages were defined as primordial stage with a full layer of squamous pre-granulosa cells, primary stage with a full layer of cuboidal granulosa cells, and secondary stage with two layers of granulosa cells. Follicles were lysed at room temperature for 5 min, centrifuged at 800 *g* for 30 s before snap-freezing in liquid nitrogen.

### Single follicle RNAseq library preparation

In total, 174 follicles (153 primordial, 41 primary, 34 secondary) were collected and processed for single follicle RNA-sequencing using the established SMART-Seq2 protocol (Trombetta et al. 2014). Briefly, cDNA was reversed transcribed from individual follicles using Maxima RT (Thermo Fisher Scientific) and whole transcriptome amplification (WTA) was performed. WTA products were purified using the Agencourt AMPure XP beads (Beckman Coulter, IN, USA) and used to prepare paired-end libraries with Nextera XT (Illumina, CA, USA). Libraries were pooled and sequenced on a NextSeq 550 sequencer (Illumina) using a 75 cycle High Output Kit (v2.5). As a secondary follicle contains many more somatic cells than a primordial and a primary follicle, libraries of secondary follicles were diluted by 2-fold to compare between follicular stages fairly.

### Single follicle RNAseq data analysis and statistics

Data processing followed a series of standard bioinformatics workflows, all performed on the Dartmouth HPC cluster (Polaris SMP). First, raw sequencing data were demultiplexed using scCloud and bcl2fastq (Illumina, v2.20), followed by alignment of the resulting FASTQ files to the reference genome using STAR Aligner (81). Gene-level count matrices were generated with HTSeq (82), and quality control metrics were extracted using Picard tools. To ensure data quality, samples with over 40% mapping to ribosomal RNA or fewer than 8 million total reads were excluded from further analysis. Batch effects from sequencing across eight batches were corrected using ComBat-seq (21), which effectively minimized intra-batch variability, as verified by hierarchical clustering analysis. Differentially expressed genes between primordial, primary, and secondary stages were identified via DESeq2 (83) with an adjusted P value cutoff of 0.05. Principal component analysis was performed using variance stabilized transformed values of gene expression data, heatmaps were generated using z-score normalized data, and violin plots were generated using normalized expression data from the DESeq2 object. Gene ontology (GO) and pathway analysis were performed using Metascape (34) on upregulated genes from primordial and primary plus secondary follicles from DESeq2. Size-based differential expression analysis was performed using the limma package in EdgeR (84). Weighted gene network coexpression analysis (WGCNA) (43) was performed on raw count data and Pearson’s correlation coefficient was calculated to associate each gene module with follicle stage. Gene set variation analysis (GSVA) (44) were performed using raw count data. GSVA pathway analysis was generated using KEGG, Reactome, and WikiPathways, while GO was performed using GO biological Processes, Molecular Function, and Cellular Component.

## Contributions

Conceptualization: TKW, AKS, YC, BAG

Analysis: HV, YC, BAG, OK

Funding: TKW, AKS, BAG, FD

Data collection: YC, DDR

Paper Writing first draft: HV, BAG, YC

Figure generation: HV, YC

Draft reviewing and editing: HV, YC, OK, DDR, AKS, FD, TKW, BAG

Supervision: TKW, BAG

## Acknowledgments

The authors acknowledge that the analysis reported in this paper was supported by the Genomics Shared Resource and Dartmouth Research Computing at Dartmouth College.

## Supporting Information

**S1 Figure. Heatmap of the Pearson’s correlation coefficient among all individual follicles after batch correction.**

**S2 Figure. Additional PCA analysis.**

**S3 Figure. Volcano and MA plots of differential expression comparisons.**

**S4 Figure. Gene ontology analysis of follicle- and oocyte-size-specific genes.**

**S5 Figure. Gene ontology analysis of follicle- and oocyte-size specific genes that overlap with oocyte and somatic cell specific genes.**

## Notes

### Competing Interest Statement

The authors declare the following competing interests: A.K.S reports compensation for consulting and/or SAB membership from Honeycomb Biotechnologies, Cellarity, Ochre Bio, Relation Therapeutics, Bio-Rad Laboratories, Passkey Therapeutics, Fog Pharma, Dahlia Biosciences, and intrECate Biotherapeutics. The other authors declare no competing interests.

